# Characterization of the microbiome of Down syndrome mouse model (Ts65Dn) in standard and high-fat diet

**DOI:** 10.1101/2020.11.16.385989

**Authors:** Ilona E. Grabowicz, Marta Fructuoso, Ilario DeToma, Mara Dierssen, Bartek Wilczyński

**Affiliations:** Faculty of Mathematics, Informatics and Mechanics, University of Warsaw, Warsaw, 02-097, Poland; Centre for Genomic Regulation (CRG), The Barcelona Institute of Science and Technology, 08003 Barcelona, Spain; University Pompeu Fabra, 08003 Barcelona, Spain; Biomedical Research Networking Center on Rare Diseases (CIBERER), Institute of Health Carlos III, Madrid, Spain

## Abstract

The intestinal microbiota is known to affect its host in numerous ways and can be altered by many factors including the host genotype and high-calorie diets. Down syndrome (DS) is a genetic neurodevelopmental disorder caused by the total or partial triplication of chromosome 21. Recently, a human study reported microbiota differences between DS and euploid humans. To further explore the differences due to the genotype, we here investigated the microbiome of trisomic mice (Ts65Dn). In trisomic mice we found a significant enrichment in abundances of bacteria: *Bacteroides ovatus, B. thetaiotaomicron*, and *Akkermansia muciniphila* - the mucus-degrading and gut-health promoting species. Since diet composition has an effect on microbiota species, we studied the effect of a high-fat diet on the observed genotypic differences. Our study provides evidence that microbiomes of trisomic mice on the control diet present more inter-individual differences than WT mice. Moreover, we observed that the high-fat diet led to increased differences between individuals and this effect was even more pronounced in the trisomic than in WT mice. We validated the results obtained with widely used 16rRNA gene sequencing with the sequencing of the total RNA.

**Highlights:**

- Down syndrome (DS) model mice faecal microbiomes are characterized by an overrepresentation of *Bacteroides ovatus, Bacteroides thetaiotaomicron*, and *Akkermansia muciniphila* species.
- DS mice are characterized by higher heterogeneity of their microbiome communities than WT mice.
- High-fat diet leads to more diverse microbiome communities between mice, especially in trisomic genotype.

## Introduction

Down syndrome (DS) is one of the most prevalent intellectual disability disorders of genetic origin. DS has an incidence of 1 in 1000 births and affects more than 5 million people worldwide (World Health Organisation report) and causes numerous developmental anomalies such as intellectual disability, facial dysmorphology, congenital defect of heart and gut, immunodeficiencies (Dierssen, 2012). With development of medicine and social care the lifespan of persons with DS is significantly prolonged in the past decades (Zigman, 2013), however, this prolongation comes together with a parallel increase of risk for age-related diseases, such as Alzheimer’s disease, increased rate of infections, hypertension and obesity (Nakamura and Tanaka, 1998; Dierssen et al. 2020). Some of the DS comorbidities, such as functional bowel disease (Gevers et al. 2014), obesity (Castaner et al. 2018), reflux oesophagitis (Cho and Blaser, 2016) or Alzheimer disease (Minter et al., 2016) were recently found to be related to certain microbiome features, it became highly interesting to characterize the DS microbiome. To date, we found only one study characterizing the gut microbiome in human DS patients (Biagi et al. 2014). To our best knowledge there is no study published describing characteristics of the microbiome of any DS mouse model. The Ts65Dn mouse model (Davisson et al., 1993) is one of the most characterized DS models, as it recapitulates most of the cognitive and neural phenotypes described in DS individuals (Gardiner, 2015) and is used for testing different therapeutics for DS or investigating the disease. In our study we attempted to characterize microbiomes of Ts65Dn mice and compare with Wild Type (WT) mice. Additionally, given that the combined prevalence of overweight and obesity in individuals with DS varies between studies from 23% to 70% (Bertapelli, 2016), and that obesity is associated with different profiles of gut microbiota, we studied the effect of High-Fat Diet (HFD) on the microbiome and behavior of Ts65Dn mice. This diet reflects the common dietary direction in the modern ‘Western’ societies and it is known to produce changes in the microbiome, lead to obesity and intestinal-inflammation (Ding et al., 2010), which are also often found in DS patients.

Recent technological advances in sequencing have led to a dramatic increase in studying characteristics of microbiomes – the collective genomes of microorganisms inhabiting their host’s surfaces (Gill et al., 2006, Holmes et al., 2012). This in turn has led to finding numerous links between different microbiomes’s properties and diverse features of its host. Majority of studies done on microbiomes so far employed use of sequencing of the PCR-amplified 16S ribosomal DNA fraction (Bashiardes et al. 2016). Although it has been widely used and became a ‘standard’ way of investigating microbiomes, it also carries several limitations such a bias towards certain microbial groups due to the used primers, detecting chimeras (Goodrich et al., 2014) or not detecting some groups at all (Jovel et al., 2016). In our study we have used the common 16S approach and validated the results using Illumina Hi-Seq sequencing of the total RNA which is devoid from the primer bias.

## Results

### Ts65Dn mice have altered microbiome composition

From the 16S sequencing, we obtained in total ~5 mln reads, within the range of 147 - 318 thousand reads per sample. Expectedly, more than 99% of the reads were of ribosomal and bacterial origin. The sequencing depth was enough to discover the majority of the bacterial species present in the samples (Suppl. Fig. 1A). The most abundant phyla and species detected were the same in both genotypes, with the most numerous ones: Bacteroidetes and Firmicutes; and species *Barnesiella viscericola* (Fig. 1A and B). The vast majority of species detected in both genotypes (with read number threshold at least 20 reads across samples) were the same - 98% of the species were common (Fig. 1C). Although the overall species abundance profiles of Ts65Dn and WT were similar, we could detect significant differences as well. Considering mice at both time points, we identified 28 species being differentially abundant between Ts65Dn and WT mice (DeSeq2, FDR corrected p<0.05), of which some (*Bacteroides ovatus*) belonged to the most abundant ones (Fig. 1D, Suppl. Table 1). Among the differentially abundant species around half of them were more abundant in Ts65Dn mice and half in WT. All of the significantly differentially abundant Bacteroides species (*B. fragilis, B. vulgatus, B. thetaiotaomicron, B. ovatus*) showed higher abundance in Ts65Dn mice.

**Fig 1.**
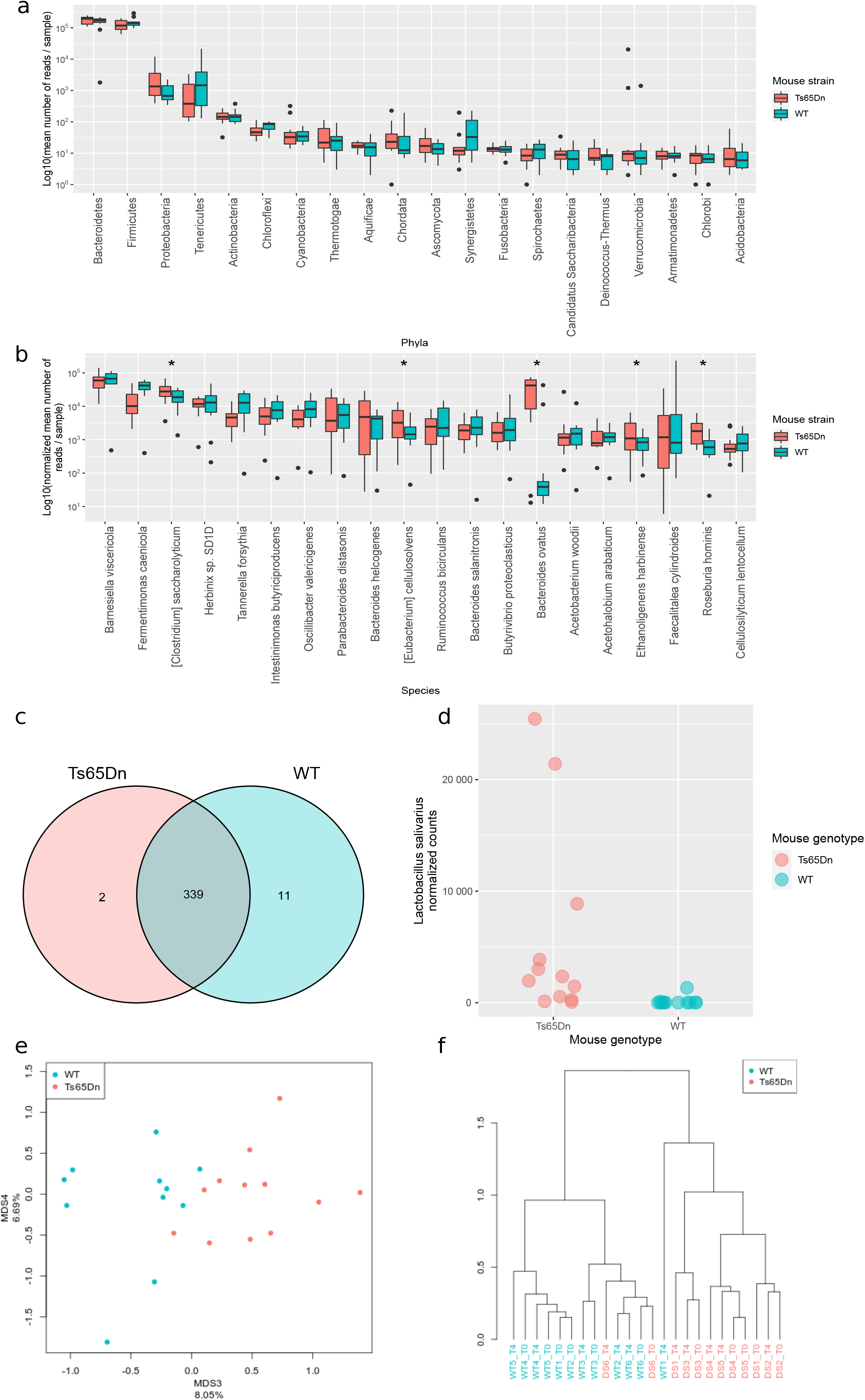
Microbiome profiles of DS and WT mice fed control and high-fat diet. A) Boxplot of the most abundant phyla detected by 16S sequencing in DS and WT mice. B) Boxplot of the most abundant species detected by 16S sequencing in DS and WT mice. C) Numbers of species detected using 16S sequencing in DS and WT mice. D) Dotplot depicting abundances of a differential species *Bacteroides ovatus* in WT and DS mice. E) Multidimensional Scaling plot of the species abundances detected in WT and DS, calculated on Canberra distance matrix F) Hierarchical clustering of samples performed on Bray-Curtis distances matrix and ward.D clustering methods

Through the multidimensional scaling analysis (MDS) we discovered that the genotype is an important factor contributing to the clustering of samples. The third leading dimension, accounting for 8% of the variance in the data, was distinguishing the Ts65Dn and WT mice groups, disregarding the diet (Fig. 1E). The first and second dimensions did not provide such separation, probably accounting for the natural diversity of the samples or noise in the data. Species mostly contributing to the separation of the two groups of samples in the leading dimension 3 in MDS were consistent with the most differentially abundant species between genotypes. Out of the top 10 species mostly contributing to the cluster separation common with the previous analysis were *B. fragilis, B. ovatus, Clostridium saccharolyticum* and *Lactobacillus salivarius*. The hierarchical clustering analysis revealed that WT and Ts65Dn mice tend to cluster according to the genotype (Fig. 1F).

### Ts65Dn mice microbiomes show a more variable response to HFD

In WT mice we found 7 microbial species with significantly changed abundance levels (DeSeq2, FDR corr. p < 0.05), while only 2 within Ts65Dn microbiomes (Suppl. Table 2 and 3). The species most significantly changing its abundance in Ts65Dn mice was *Eubacterium eligens* (corr. p = 0.04), which was the second mostly changed species also in WT mice (corr. p = 0.0001) - in both cases its abundance was decreased upon HFD. The second significantly differentially abundant species in DS mice was *Synechococcus sp. JA-3-3Ab*, however, its abundance was very low (with mean counts per Ts65Dn sample = 30). On the other hand, the most significantly changed species in WT mice - *Clostridium perfringens* - was increased significantly on HFD both in WT (corr. p = 0.0001, mean counts in WT = 237), and in Ts65Dn mice (corr. p = 0.07), though much less abundant (mean counts in Ts65Dn mice = 26). On the phylum level, we detected significant differences in abundances of Verrucomicrobia and Proteobacteria, which were overrepresented in Ts65Dn mice. The beta-diversity was significantly lower in WT mice meaning that WT individuals were characterised by their microbiome contents more similar to each other than in Ts65Dn (Fig. 2A; Wilcoxon test on correlation values, W = 640, p = 0.005), (WT mice correlations: median Pearson R = 0.81, p = 1.75 e^-80^; Ts65Dn mice correlations: median Pearson R = 0.60, p = 4.67 e^-34^). The HFD itself increased the beta-diversity in both WT and Ts65Dn mice, as the correlations of species abundances between the samples were decreasing (Pearson R medians 0.69 to 0.57 and 0.83 to 0.67 in Ts65Dn and WT mice, respectively, Fig. 2A). To compare these results with humans we re-analysed the Biagi et al. (2014) data coming from healthy humans and humans with DS. In humans we observed a lower number of species present, however, there were 252 species in common with mice (Fig. 2B). Moreover, we observed significantly reduced within-group correlations between humans than in mice (humans: median Pearson R = 0.5, p = 1.5e^-7^; mice: median Pearson R = 0.72, p = 5.7e^-17^; Wilcoxon test W = 23583, p-value = 1.7-10, Fig. 2C). This result is not surprising, as mice were grown in the controlled conditions and they were similar genotypically to each other, which was not the case in non-related humans. However, unlike in mice, we observed lower inter-individual correlations in euploid humans than in DS individuals (Fig. 2C). This means that the microbiome profiles were more similar to each other in DS individuals than in euploid, healthy humans.

**Fig 2.**
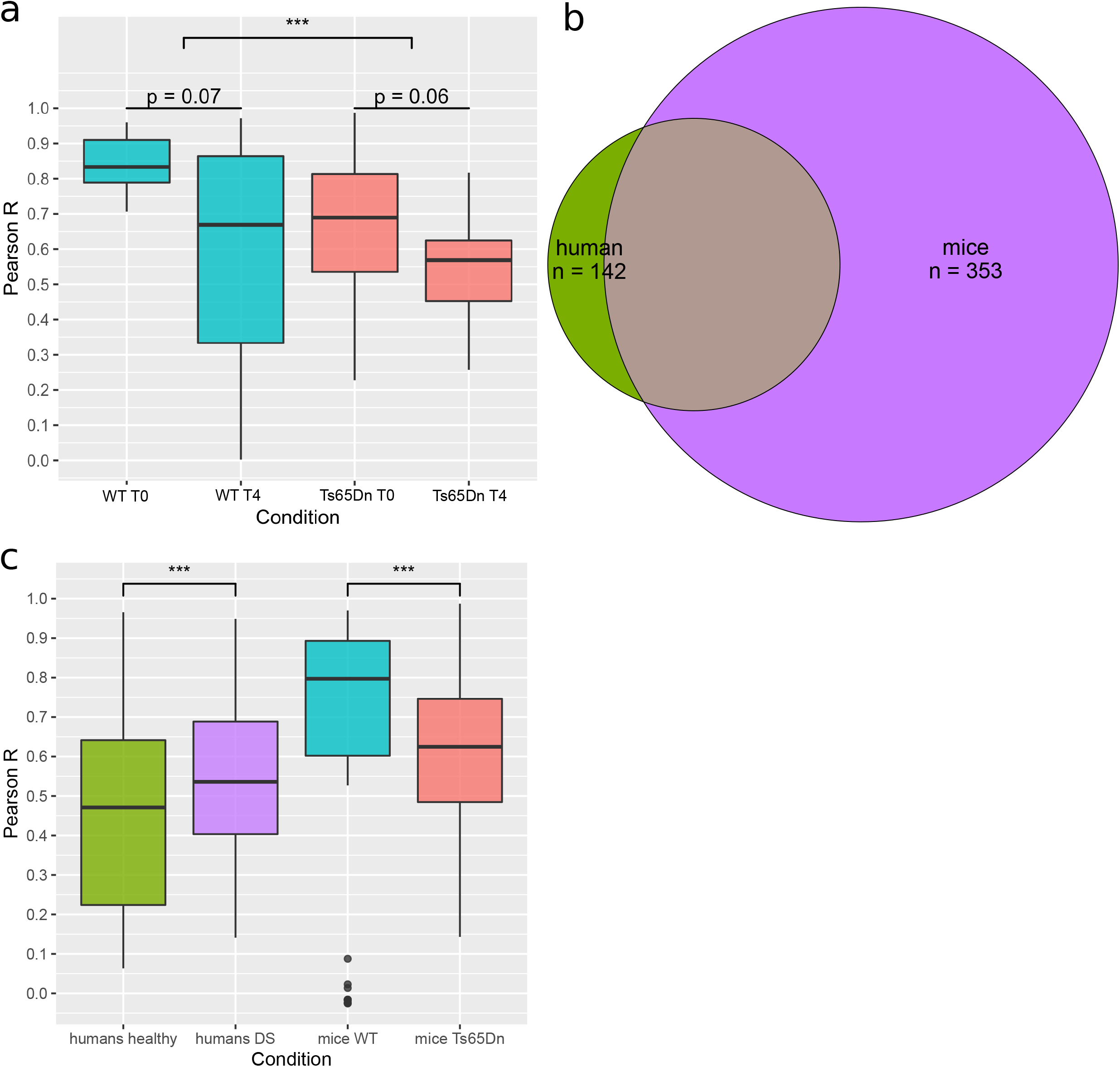
Individuals affected by DS exhibit higher inter-individual (beta-)diversity. A) Boxplots of Pearson correlation R values. Correlations were calculated between the samples belonging to each of the groups with different mouse strains and HFD time points. B) Venn diagram showing numbers of common and different species between humans and mice (healthy and with DS, collectively). C) Boxplots of Pearson correlation R values. Correlations were calculated between the samples belonging to each of the groups: healthy humans, humans with DS, WT mice, DS mice. Human data were re-analyzed and come from Biagi et al. 2014.

### Results validation using total RNA sequencing

To validate the results obtained with sequencing of the 16s rRNA gene (Illumina HiSeq), we also used total RNA sequencing, which consists mainly of rRNA. Due to a lack of samples from the same time points, we used samples coming from mice after 2 and 14 days of the introduction of HFD of WT and Ts65Dn mice. From the total RNA sequencing, we obtained in total ~115 mln reads, within the range of 11 - 15 10^6^ reads per sample. Expectedly, more than 94% of the reads were of ribosomal origin. After filtering only for the ribosomal reads, 91% of them were assigned to bacterial species. The sequencing depth was enough to discover the majority of the bacterial species present in the samples (Suppl. Fig. 1B). All the species detected by 16S sequencing were also discovered in the total RNA sequencing (Fig. 3A, Suppl. Fig. 1C (WT mice), Suppl. Fig. 1D (Ts65Dn mice)). To verify whether for the species detected in both types of sequencing their abundances were concordant, we calculated the correlation coefficients for the mean species abundances in both sequencing modalities. The high correlation (Pearson R = 0.67, p = 1.43 e^-46^) suggests that both methods give similar results for the species that can be detected by both methods (Fig. 3B). Among differentially abundant species between mouse strains detected by total RNA sequencing, 3 were in common with 16S sequencing: *Bacteroides ovatus, Bacteroides thetaiotaomicron*, and *Akkermansia muciniphila* (Deseq2, FDR p < 0.05). Concordantly between methods, all 3 species were found to be overrepresented in trisomic mice.

In order to validate the similarity of the microbial profiles among the samples coming from the same mouse strain, we calculated the correlation coefficients for all the WT or Ts65Dn samples (disregarding the time point to obtain more samples in each group) also in the total RNA dataset. Median R values for correlations of the species abundances between samples were again lower in trisomic than WT mice (Suppl. Fig. 1E; median R = 0.91, p = 0 and R = 0.82, p = 0, for WT and Ts65Dn mice, respectively), however, the difference did not reach statistical significance, probably due to low sample size (Wilcoxon test, W = 40, p = 0.31).

**Fig 3.**
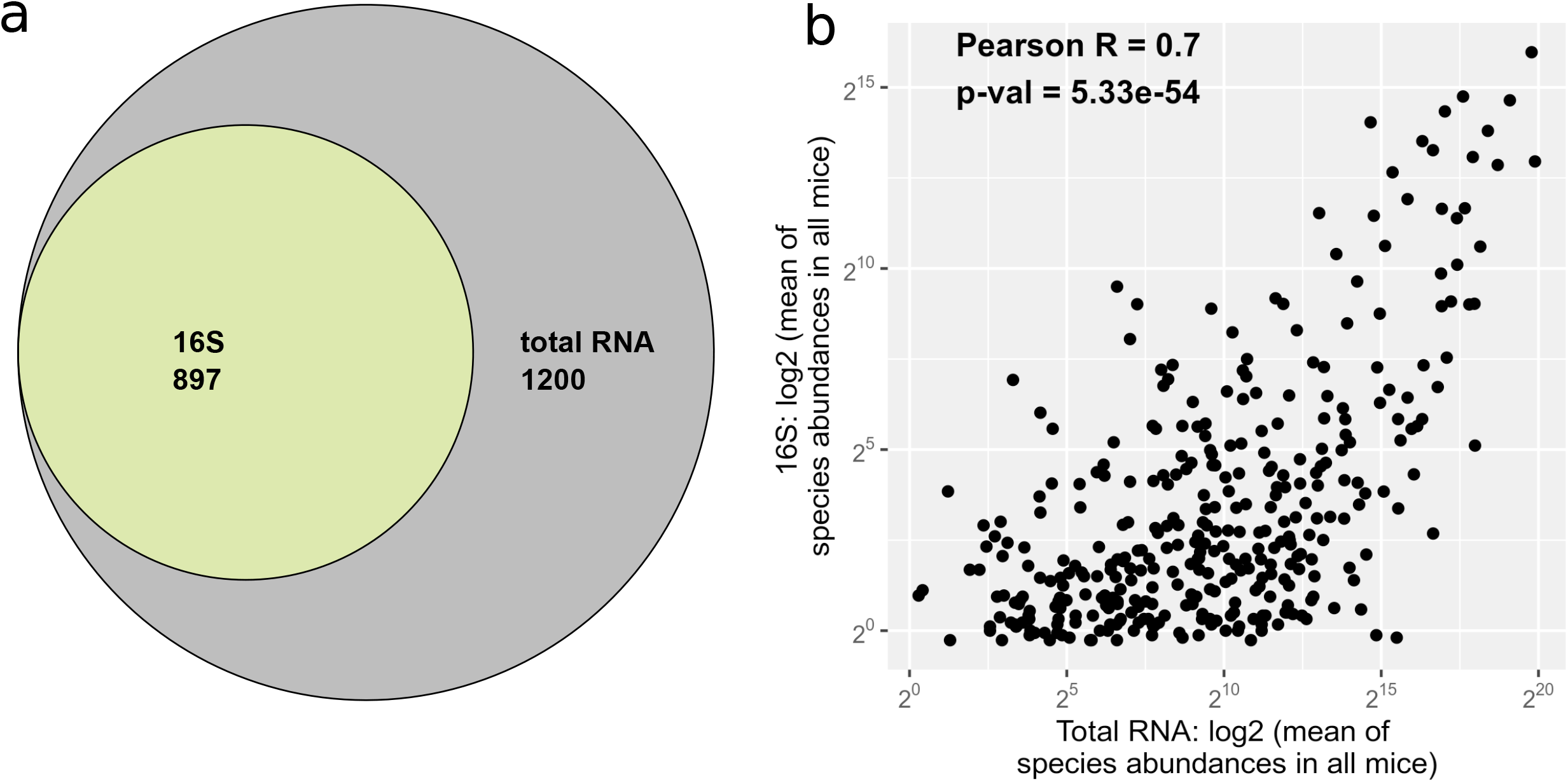
Validation of 16S results by total RNA sequencing. A) Venn diagram depicting the overlap between numbers of species detected using 16S and total RNA sequencing B) Dotplot showing log2 values of species abundances detected by 16S and total RNA sequencing. Dots are shown only for the species which were detected by both methods. Each dot represents each species.

## Materials and Methods

### Mice, housing and sample collection

5 months-old male wild-type (WT) and Ts(1716)65Dn (Ts65Dn) mice were obtained through crossings of a B6EiC3Sn a/A-Ts(1716)65Dn female to B6C3F1/J males purchased from The Jackson Laboratory (Bar Harbor, USA). Genotyping was performed by amplifying genomic DNA obtained from the mice tail (Liu et al. 2003). The colony of Ts65Dn mice was bred in the Animal Facility of the Barcelona Biomedical Research Park (PRBB, Barcelona, Spain), and until the experiments, mice received chow and water ad libitum in controlled laboratory conditions with the temperature maintained at 22 ± 1 degree C and humidity at 55 ± 10% on a 12h light/dark cycle (lights off 20:00h). All experimental procedures were approved by the local ethical committee (Comité Ético de Experimentación Animal del PRBB (CEEA-PRBB); procedure number MDS-12-1464P3), and met the guidelines of the local (law 32/2007) and European regulations (EU directive n° 86/609, EU decree 2001-486) and the Standards for Use of Laboratory Animals n° A5388-01 (NIH).

### Experimental groups and diets

All mice were individually housed in multi-take metabolism cages (PheCOMP, Panlab-Harvard Instruments, Barcelona, Spain) and provided with standard chow (SC) and water for two weeks to allow habituation to the housing conditions. After habituation, mice were assigned to each diet-condition. These mice served for the assessment of the genotype-dependent differences in feeding behavior and microbiome composition in a nutritionally balanced diet condition.

To investigate the effect of obesogenic high-fat diet (HFD) of microbiome composition in trisomic mice, all mice were fed free-choice HFD and faecal samples were collected before the introduction of HFD and after 2, 14 and 28 days of HFD. To obtain information about the gut microbial profile of trisomic mice we sequenced the: 16S rRNA gene sequences from 6 Ts65Dn and 6 WT mice; and the total RNA from 2 Ts65Dn and 3 WT mice, to validate the 16S results. The mice fed HFD had free access to HFD pellets (HFD, Test Diet, USA) and SC (SC SDS, UK). For the 16S rRNA sequencing the faecal samples were collected under standard chow diet (SC SDSand after 28 days of HFD, while for the total RNA sequencing after 2 and 14 days of HFD (day 14: WT n=2). Pellets were renewed at least twice a week to ensure the maintenance of their organoleptic properties.

### 16S sequencing

DNA was extracted from faecal samples from Ts65Dn (n = 6) and WT (n = 6) mice on day 0 (SC) and 28 (28th day of high-fat diet). Libraries were prepared using Illumina protocol targeting the V3 and V4 variable regions of the 16S rRNA gene. This protocol involves a two-step tailed PCR approach that generates ready-to-pool amplicon libraries. The protocol includes the primer pair sequences for the V3 and V4 regions that create a single amplicon and overhang adapter sequences that must be appended to the primer pair sequences for compatibility with Illumina index and sequencing adapters. During the first PCR, locus-specific primers contained sequence tails that allowed for a second PCR to add Nextera^®^ XT indexed adapters. The PCR was performed in 10 μl volume with 0.2 μM primer concentration. Cycling conditions were: Initial denaturation of 3 min at 95 °C followed by 25 cycles of 95 °C for 30 sec, 55 °C for 30 sec, and 72 °C for 30 sec, ending with a final elongation step of 5 min at 72 °C. As a control for downstream procedures, we also used two DNA samples derived from bacterial mock communities obtained from the BEI Resources of the Human Microbiome Project.

The overhang adapter sequence must be added to the locus - specific primer for the region to be targeted. After the second PCR 25 μl of the final product was used for purification and normalization with SequalPrep normalization kit (Invitrogen), according to the manufacturer’s protocol. Sequencing was performed using Illumina MiSeq with 2×300 bp reads using v3 chemistry with a loading concentration of 10 pM. In all cases, 15% of PhIX control libraries was spiked in, to increase the diversity of the sequenced sample.

### Total RNA sequencing

Total RNA was extracted from faecal samples from Ts65Dn (n=2) and WT (n=3) mice fed with HFD collected on day 2 and day 14 of the experiment (day 14: WT n=2). Libraries were prepared using NEBNext Ultra Directional RNA Library Prep Kit for Illumina (New England Biolabs). Validation and quantification of libraries were done using Agilent 2100 Bioanalyzer with Agilent RNA 6000 Pico Kit. Sequencing was performed using 2×100 bp paired-end mode on HiSeq 4000 (Illumina) in Genomed SA (Warsaw, Poland) according to the manufacturer’s protocol.

### Data processing

The quality of all the sequences was determined using FastQC (Andrews, 2010). In the case of 16S sequencing reads quality of the data was good. The reverse reads were of worse quality at the 3’ ends, therefore they were subjected to trimming by 50 nucleotides (Trimmomatic, Bolger at el. 2014). RNA HiSeq data, quality of all reads was excellent – it was falling in the ‘green’ confidence area shown by FastQC. Share of ribosomal reads was assessed with SortMeRNA tool (Kopylova et al. 2012) using the default settings and all of the available reference databases. Sequences were then mapped to microbial taxa using seed-kraken (Brinda et al. 2015, Wood and Saltzberg 2014). The database for seed-kraken was built based on a collection of sequences from RefSeq (Feb. 2016): Bacteria, Viruses, Fungi, H. sapiens, and M. musculus. Sequences needed to have version_status==‘latest’ and assembly_level = “Complete Genome” or assembly_level = “Chromosome”.

After mapping reads to taxa we have split them into different taxonomic levels. After excluding the unclassified and mouse reads, the remaining ones were split into taxonomic levels and normalized (normalization to the same total number of reads in each mouse).

### Bioinformatic and statistical analysis

All analyses were done using scripts written in python or R programming languages. Venn diagrams were produced using an R package ‘venn’ (Adrian Dusa). Rarefaction curves depict numbers of observed species and were generated with a self-written script in R. The negative binomial tests were performed using ‘DeSeq2’ (Love et al. 2014) with the default settings. Hierarchical Clustering was done in R using package ‘vegan’ (Oksanen et al. 2007) and illustrated using package ‘dendextend’ (Galili, 2015). Multidimensional Scaling was done using the package ‘vegan’ (Oksanen et al. 2007). All the other illustrations were generated also in R (‘ggplot2’ (Wickham 2009), ‘ggrepel’ (Slowikowski 2016), ‘randomcoloR’). Other R packages used to manipulate the data: ‘plyr’ (Hadley Wickham), ‘data.table’ (Matt Dowle), ‘scales’, ‘dplyr’, ‘reshape2’, ‘grid’). The bash, python, and R code to reproduce the results is available at: https://github.com/ilona-grabowicz/microbiome-in-DS

### Data availability

The data for this study have been deposited in the European Nucleotide Archive (ENA) at EMBL-EBI under accession number PRJEB40927 and PRJEB41352. (https://www.ebi.ac.uk/ena/browser/view/PRJEB40927 https://www.ebi.ac.uk/ena/browser/view/PRJEB41352).

## Discussion

In our study, we examined differences in microbiome content between the Ts65Dn trisomic mice, a validated DS model and their WT littermates. The key finding of our study is that we identified 3 bacterial species significantly overrepresented in Ts65Dn mice using two sequencing methods (16S rRNA gene and total RNA): *Bacteroides ovatus, Bacteroides thetaiotaomicron*, and *Akkermansia muciniphila. B. fragilis* and *B. vulgatus* were also more abundant in Ts65Dn mice as discovered by 16S and total RNA sequencing, but only reached significance in 16S. All those species (*B. ovatus, B. thetaiotaomicron, B. fragilis, B. vulgatus* and *A. muciniphila*) are capable of harvesting host mucus glycans (Marcobal et al. 2011, Dao et al. 2016), which would suggest a possible mucus overproduction in DS mice. Previous studies showed proliferation impairment in intestinal germinative niches, which differentiate into the goblet cells that secrete mucus (Fuchs et al. 2004). This precursor proliferation reduction could thus be to some extent aggravated by the microbial composition in Ts65Dn mice and open therapeutic possibilities for DS abnormal mucosa formation and secretion/absorption problems.

Moreover, within Ts65Dn mice we found a lower bacterial alpha-diversity (less species found), which has also been described in obese individuals (Liu et al. 2017, LeChatelier et al. 2013, Cotillard et al. 2013) along with higher differences between samples (beta-diversity) confirmed by two sequencing methods, indicating a more heterogeneous community structure within obese individuals (Liu et al. 2017).

Numerous studies have demonstrated a link between microbiota composition and its host behavior (Sampson and Mazmanian 2015). Interestingly, *B. ovatus* abundance is significantly higher in Attention-Deficit/Hyperactivity Disorder (ADHD) patients and has been associated with ADHD symptoms (Wang et al. 2019). Among children with DS, the prevalence of ADHD was reported to be very high, around 43% (Ekstein et al. 2011), and could be also related with the hyperactivity and the reduced attention detected in Ts65Dn mice (Escorihuela et al. 1995, Whitney and Galen, 2013, Driscoll et al. 2004).

Given the obesity comorbidity in DS, we studied how the microbiome profile of Ts65Dn changed upon high-fat diet feeding. In standard chow conditions, Ts65Dn are in fact leaner than their wild type littermates (Fructuoso et al., 2018; Dierssen et al. 2020), and we found *A. muciniphila* overrepresentation in Ts65Dn, which is associated with lowering body fat mass (Everard et al. 2013) and improving glucose homeostasis (Shin et al. 2014). Interestingly, upon access to a high-fat diet Ts65Dn mice present increased body adiposity and gain more weight than WT (Fructuoso et al. 2018). The higher abundance of *A. muciniphila* has also been associated with lower leptin levels and lower adipocyte diameter (Dao et al. 2016). Leptin is a hormone produced predominantly by adipocytes and enterocytes and it inhibits the feeling of hunger (Brennan and Mantzoros, 2006). Interestingly, Everard et al (2013) found lower *A. muciniphila* abundance in leptin-deficient obese mice, and we previously demonstrated that Ts65Dn mice present increased leptin levels compared to WT (Fructuoso et al., 2018), but consumed more calories, suggesting that leptin would be ineffective in controlling satiety in DS. In humans results are divergent as in one study the higher *Akkermansia* abundance was found in subjects with normal glucose tolerance (Zhang et al. 2013) while in another one in type2 diabetic patients (Qin et al. 2012). Also in individuals with DS, while leptin levels in plasma correlate with adiposity as in the general population, they present lower (Radunovic et al., 2003, Yahia et al., 2012, Gutierrez-Hervas et al., 2020) or higher (Proto et al., 2007; Magge et al., 2008) leptin levels as compared to matched controls without DS.

Moreover, Liu et al. (2017) found enrichment in *Bacteroides ovatus* and *B. thetaiotaomicron* as well as *A. muciniphila* in lean controls compared to obese mice. All these three species were enriched in the trisomic mice, which at baseline are leaner but presented higher body weight increase upon HFD feeding. In fact, the same study indicated that in HFD fed mice, gavage with live *B. thetaiotaomicron* bacteria resulted in a concomitant increase of *A. muciniphila*, indicating that these bacterial strains might have positive interactions, what could also explain why we found their joint increase.

To the best of our knowledge, our study is the first that describes the microbiome profiles of a DS mouse model. In humans, there is only one study (Biagi et al. 2014) describing the microbiomes of individuals with DS. The bacterial families for which DS individuals were enriched in that study were Parasporobacterium and Sutterella, which we did not find in the mouse microbiome (neither sequenced with 16S nor total RNA). Sutterella has been reported to present exclusively in the human gut (Krych et al. 2013). The family *Veillonellaceae*, was significantly decreased in individuals with DS and was also decreased in Ts65Dn mice (in total RNA sequencing), although the difference did not reach statistical significance. On the other hand, we found a significant decrease in DS mice of the genus *Veillonella* (16S sequencing). This difference might result from the natural differences occurring between human and murine microbiomes.

In conclusion our results revealed genotype- and diet-dependent differences in the microbiome profile of trisomic Ts65Dn that could explain some DS behavioral and metabolic phenotypes. Further studies are granted to get deeper insight into mechanism and to propose microbiome-based therapy as an option for DS. Additionally, we provided evidence that sequencing the total RNA provides an interesting alternative to sequencing of the 16 rRNA gene as it allowed to identify much more bacterial species and it’s not biased by the selection of primers for PCR.

---

**Suppl. Fig. 1.**
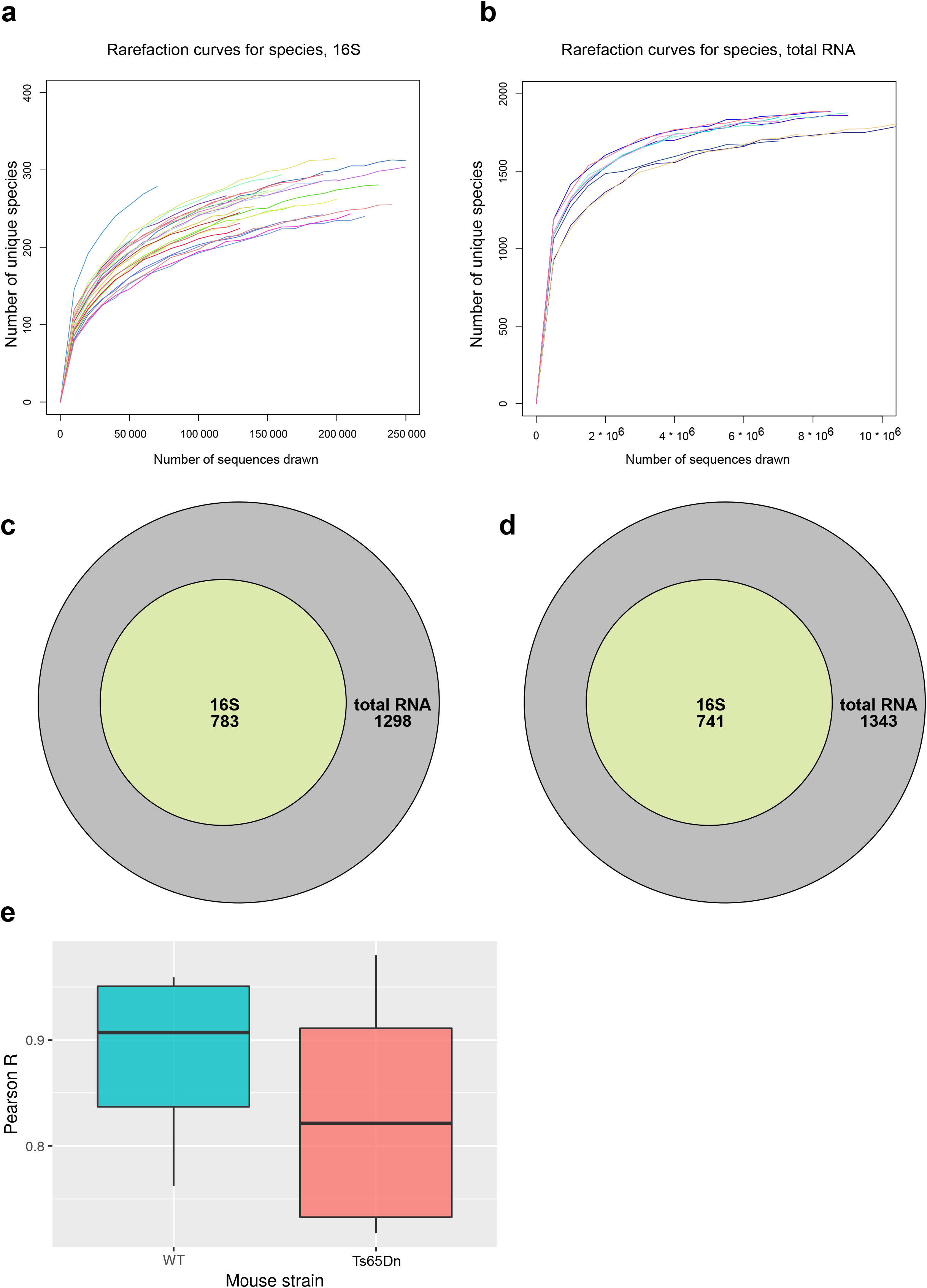
Comparison of results obtained by 16S sequencing with total RNA sequencing. A) Rarefaction plot for the species detected by 16S rRNA gene sequencing. Each color depicts one faecal sample. B) Rarefaction plot for the species detected by total RNA sequencing. Each color depicts one faecal sample. C) Venn diagram of the species present in WT mice detected by total RNA sequencing (Illumina HiSeq) D) Venn diagram of the species present in DS mice detected by total RNA sequencing (Illumina HiSeq) E) Boxplot of Pearson correlation levels between samples coming from the same mouse strain, detected by total RNA sequencing (Illumina HiSeq).

## Acknowledgements

The authors thank Pedro Gonzalez Torres and Toni Gabaldon for help with sequencing. IEG and BW have been supported by Polish National Center for Research and Development grant [ERA-NET-NEURON/10/2013] and Polish National Science Centre grant [DEC-2015/16/W/NZ2/00314]. The laboratory of M. Dierssen is supported by DIUE de la Generalitat de Catalunya (Grups consolidats 2017SGR926, 2017SGR595). MD also acknowledges the support of Spanish Agencia Estatal de Investigación (PID2019-110755RB-I00/ AEI / 10.13039/501100011033; ERA-NET-NEURON/10/2013 and EMBL partnership), the Centro de Excelencia Severo Ochoa, CERCA Programme/Generalitat de Catalunya, H2020 SC1 Gene overdosage and comorbidities during the early lifetime in Down Syndrome GO-DS21-848077, Fundació La Marató De TV3 (201620-31_MDierssen), EU (JPND HEROES), NIH (Grant Number: 1R01EB 028159-01). MF, IDT and MD were supported by Fondation Jérôme Lejeune (2019b - Project #1887) grant.

## Authors contributions

IEG and BW conceived and designed the study. IEG performed the bioinformatics and statistical analyses, interpreted and visualized results and wrote the original draft; MF and MD organised and performed the behavioral experiments. MF collected and prepared the samples. IDT commented on the manuscript. All authors discussed the results and reviewed and edited the manuscript.

